# No evidence for schistosome parasite fitness trade-offs in the intermediate and definitive host

**DOI:** 10.1101/2022.11.11.516207

**Authors:** Winka Le Clec’h, Frédéric D. Chevalier, Kathrin Jutzeler, Timothy J.C. Anderson

**Affiliations:** Host parasite Interaction program, Texas Biomedical Research Institute, P.O. Box 760549, 78245 San Antonio, Texas, USA; UT Health, Microbiology, Immunology & Molecular Genetics, San Antonio, TX 78229; Disease Intervention and Prevention program, Texas Biomedical Research Institute, P.O. Box 760549, 78245 San Antonio, Texas, USA

**Keywords:** *Schistosoma* parasite, *Biomphalaria* snail host, rodent host, virulence-transmission trade-offs, positive pleiotropy, genetic crosses, selection

## Abstract

**Background:** The trematode parasite *Schistosoma mansoni* uses an aquatic snail intermediate and a vertebrate definitive host to complete its lifecycle. We previously showed that a key transmission trait – the number of cercariae larvae shed from infected *Biomphalaria spp*. snails – varies significantly within and between different parasite populations and is genetically controlled by five loci. We investigated the hypothesis that the success of parasite genotypes showing high fitness in the intermediate snail host, may be offset by lower fitness in the definitive vertebrate host.

**Methods:** We investigated this trade-off hypothesis by selecting parasite progeny producing high or low number of larvae in the snail, and then comparing fitness parameters and virulence in the rodent host. We infected inbred BALB/c mice using two *Schistosoma mansoni* parasite lines (high shedder (HS) and low shedder (LS) lines), isolated from F2 progeny generated by genetic crosses between SmLE (HS parent) and SmBRE (LS parent) parasites. We used the F3 progeny to infect two populations of inbred *Biomphalaria glabrata* snails. We then compared life history traits and virulence of these two selected parasite lines in the rodent host to understand pleiotropic effects of genes determining cercarial shedding in parasites infecting the definitive host.

**Results:** HS parasites shed high numbers of cercariae, which had a detrimental impact on snail physiology (measured by laccase-like activity and hemoglobin rate), regardless of the snail genetic background. In contrast, selected LS parasites shed fewer cercariae and had a lower impact on snail physiology. Similarly, HS worms have a higher reproductive fitness and produced more viable F3 miracidia larvae than LS parasites. This increase in transmission is correlated with an increase in virulence toward the rodent host, characterized by stronger hepato-splenomegaly and hepatic fibrosis.

**Conclusions:** These experiments revealed that schistosome parasite fitness was positively correlated in intermediate and definitive host (positive pleiotropy). Therefore, we rejected our trade-off hypothesis. We also show that our selected schistosome lines exhibit low and high shedding phenotype regardless of the intermediate snail host genetic background.

## BACKGROUND

Organisms with complex lifecycles, such as parasites, face a central issue: genetic variants that increase fitness in one lifecycle stage may be deleterious in other stages. However, polymorphisms in life-history strategies exist and parasite populations can exhibit striking differences in their transmission/virulence trade-offs and adopt different strategies. For example, in the cestode *Schistocephalus solidus*, different parasite populations exhibit large differences in virulence toward their stickleback fish host (1–3).

We encountered a similar case with two *Schistosoma mansoni* parasite populations that differ in the amount of larvae produced (4). Schistosome parasites have a complex lifecycle, involving a freshwater intermediate snail host and a mammal definitive host. Larvae penetrate the snail head-foot, differentiate into sporocysts that then asexually proliferate to generate daughter sporocysts. The daughter sporocysts release cercariae, the mammal-infective larval stage of the parasite. Hundreds to thousands of these motile cercariae exit through the snail body wall and are released into freshwater. Exit through the body wall results in leakage of hemolymph and damage to the snail. Motile cercariae larvae actively locate and penetrate mammal host skin, where they mature into adult worms and reproduce, starting a new cycle. Of the two Brazilian *S. mansoni* parasite populations studied here, one (referred as “high shedder” or HS) develops more sporocyst cells and produces larger numbers of cercariae, but causes rapid mortality of infected snails. The second population (referred as “low shedder” or LS) however, develops less sporocyst cells, and produces fewer cercariae, resulting in lower snail mortality (4).

Interestingly, Gower and Webster (5) have shown that pathogen fitness in the intermediate snail host was inversely correlated with pathogen fitness in the definitive host in a different laboratory schistosome population. Such antagonistic pleiotropy can promote balanced polymorphisms (6,7). While antagonistic pleiotropy is considered a minor contributor to balancing selection (8), it is involved in the maintenance of phenotypic variation in both animals [e.g. horn size in Soay sheep through a trade-off between survival and reproductive success (9), or adult size and egg-to-adult development in seaweed fly *Coelopa frigida* (10)] and plants [e.g. flower size in yellow monkeyflower through a trade-off between viability and fecundity (11–13), or in temporal variation of pollen competitive ability in maize (14)]. However, antagonistic pleiotropy is not ubiquitous: in *Drosophila*, the lack of genetic trade-offs between lifespan and fertility or other life history traits like resistance to oxidative chemical in long-lived populations have shown instead clear evidence for positive pleiotropy (15).

To test which pleiotropic effect plays in our parasite-hosts system, we measured transmission stage production, as well as virulence (measured as the parasite-induced rodent morbidity and the alteration of snail physiological parameters), of our schistosome populations on rodent definitive host and in different populations of intermediate snail hosts. Using genetic crosses between LS and HS *S. mansoni* parasite populations combined with classical linkage mapping, we previously demonstrated that cercariae production trait is polygenic, with additive variation at 5 different QTLs (quantitative trait loci) explaining 28.56% of the variation in cercarial production (16). Taking advantage of these genetic crosses, we selected F2 progeny exhibiting low (LS) or high (HS) cercarial shedding profiles, to generate F3 populations with extreme transmission phenotypes (Figure 1).

**Figure 1:**
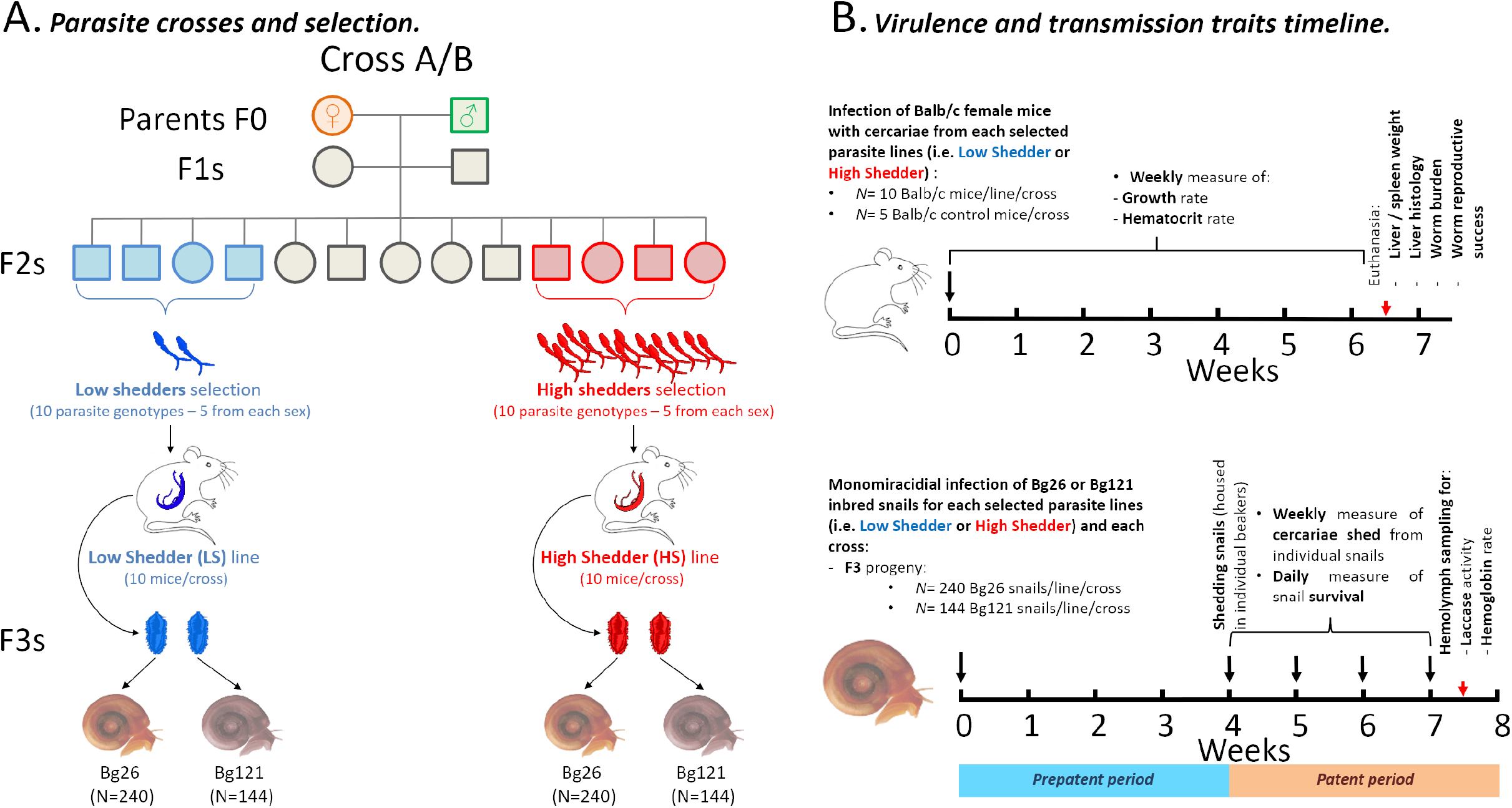
Outline of the parasite genetic crosses and the timeline of life-history traits measured in hosts. **(A)** We performed two independent reciprocal genetic crosses between single genotypes of SmLE-HS and SmBRE-LS *Schistosoma mansoni* parasite populations exhibiting striking differences in term of virulence and transmission stage production (number of cercariae produced) (16, 23). For both crosses, we generated F3 progeny by selecting F2 parasites exhibiting both extreme shedding phenotype. **(B)** We measured fitness and transmission traits of HS and LS parasites in the rodent host (BALB/c female mice) during the infection (growth and hematocrit rate) and at euthanasia (parasitic worm burden, parasite reproductive fitness, mouse spleen and liver weight, and liver histopathology). We also measured fitness and transmission phenotypes of HS and LS parasites infecting two different *Biomphalaria glabrata* inbred snail (Bg26 and Bg121), during the patent period of the infection (survival and cercarial production). We evaluated the virulence of these parasite lines by measuring the snail survival rate, the total laccase-like activity, as well as the hemoglobin rate in infected snail hemolymph samples.

This study examined the transmission/virulence trait and potential trade-off in the selected third-generation progeny (F3) on the mammal definitive host (BALB/c mice) and on two different inbred populations of snails. We demonstrated that LS parasites show reduced reproductive fitness, associated with low production of miracidia or cercariae and less damage to both rodent or snail hosts. In contrast, HS parasites show higher reproductive fitness, resulting in more larvae produced and linked to a higher morbidity of their rodent and snail hosts. These results do not reveal fitness trade-offs. Instead we observed positive pleiotropy in our schistosome-snail-rodent model.

## METHODS

### Ethics statement

This study was performed in accordance with the Guide for the Care and Use of Laboratory Animals of the National Institutes of Health. The protocol was approved by the Institutional Animal Care and Use Committee of Texas Biomedical Research Institute (permit number: 1420-MU).

### Overview of study design

Our study design is summarized in Figure 1, and methodology of each stage is explained below.

### Biomphalaria glabrata *snails and* Schistosoma mansoni *parasites*

Uninfected inbred albino *Biomphalaria glabrata* snails (line Bg26 and line Bg121, both derived from 13-16-R1 line; (17) were reared in 10-gallon aquaria containing aerated freshwater at 26-28°C on a 12L-12D photocycle and fed *ad libitum* on green leaf lettuce. All snails used in this study had a shell diameter between 8 and 10 mm, as snail size can influence cercarial production (18, 19). We used inbred snails to minimize the impact of snail host genetic background on the parasite life history traits (4).

The low shedder (LS) and high shedder (HS) *S. mansoni* lines were generated by selection of F2 progeny exhibiting either low or high production of cercariae. These F2s were generated by two independent genetic crosses of single genotypes from the SmLE-H population (high shedder parent) and the SmBRE-L population (low shedder parent) (Figure 1) (16).

### *Selection of low (LS) and high (HS) shedding F2* S. mansoni *progeny and mice infection*

For each cross (cross A and cross B) conducted by (16), we measured cercarial production of individual snails infected with F2 parasite progeny (N=204 F2A and N=204 F2B) over the 4 weeks of the patent period (4-7 weeks post infection). Each snail was infected with a single miracidia and isolated in a uniquely labeled 100 mL glass beaker filled with ^~^50 mL freshwater at the first shedding. Snails were fed *ad libitum* with fresh lettuce and kept in the dark in a 26-28°C temperature-controlled room.

At week 6, we identified 30 infected snails from each cross producing the lowest average number of cercariae (N=30 F2A-Low and N=30 F2B-Low) and 30 infected snails producing the highest average number of cercariae (N=30 F2A-High and N=30 F2B-High). We determined parasite gender by PCR amplification of specific sex markers on gDNA extracted from clonal cercariae (20) collected from each of these selected infected snails.

For each cross, we generated two lines of parasites from the F2s: a low shedder (LS) line and a high shedder line (HS). For each line, five female and five male parasite genotypes were selected for producing the F3 generation of parasites (Figure 1).

At week 7, we infected 10 female BALB/c mice (8 weeks old) per line (LS and HS) and per cross (A and B) with 80 cercariae of the female genotypes (16 cercariae from 5 snails each) and 80 cercariae of the male genotype (16 cercariae from 5 snails each). Cercariae of each gender were counted under a microscope, transferred into a glass vial filled with freshwater and uniquely labelled mice were infected by tail immersion (21). Control mice (N=10 - 5 per cross) were treated the same way but tails were immersed in freshwater only, without cercariae.

After 44 days, we euthanized and perfused each mouse to recover and count the F2 adult male and female parasite worms for each line (LS and HS) and cross (A and B). We also collected mouse livers containing the F3 eggs (Figure 1).

### *Measurement of* S. mansoni *pathogenicity in mouse definitive host*

#### a. Mouse physiological response to parasitic infection

##### i. Growth rate

We weighted each mouse weekly using a precision scale (Metler Toledo). We recorded the growth rate of each mouse as the ratio between the weight at *t+x* days after infection and the weight at *t0* (day of infection with LS and HS parasite lines) as follows:

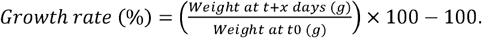

##### ii. Hematocrit rate

We collected blood from each mouse weekly, by tail-tip amputation, using microhematocrit capillary tubes (10 μL) and measured the hematocrit rate for each mouse using a Critocaps Micro-Hematocrit Capillary Tube Reader (Leica Microsystems) (22).

#### b. Mouse pathology: liver and spleen

##### i. Liver and spleen weight

We euthanized and dissected mice (t+44 days after infection) and weighed livers and spleens. *S. mansoni* infection causes hepatomegaly (inflammation of the liver due to eggs trapped in hepatic tissue) and splenomegaly (enlargement of the spleen, due to the inflammatory response to the parasitic infection) and therefore, the ratio between liver weight/mouse body weight or spleen weight/mouse body weight serve as proxies to evaluate the pathogenicity of the selected *S. mansoni* lines on the rodent host.

We recorded the ratio liver or spleen weight/mouse body weight for each mouse as follows:

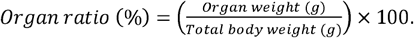

##### ii. Liver histology: egg number, granuloma size, and hepatic fibrosis

We stored the right frontal lobe of each liver in 10% neutral buffered formalin (N=50 liver samples) for further processing and embedding into paraffin. Each tissue block was cut at a thickness of 4 μm using a Microm HM325 rotary microtome. We obtained twelve sections at 60 μm intervals which were mounted onto four slides per liver. The slides were alternately stained with hematoxylin-eosin for quantitative analysis of granulomas or Mason’s trichrome to detect collagen deposition and to quantify fibrosis due to lodged parasite eggs. All slides were scanned with the Zeiss Axio Scan.Z1 whole slide scanner at a resolution of 0.22 μm/pixel. Using HALO^®^ software (v3.4, Indica labs), we analyzed sections that were at least 120 μm apart. We counted the number of eggs lodged in liver tissue and calculated egg density by dividing the number of eggs by the analyzed tissue area. We annotated and quantified the area of individual granulomas (5-17/liver) surrounding a single parasite egg. We recorded 209 granulomas for HS lines (cross A: N = 111, cross B: N = 98) and 181 for LS lines (cross A: N = 94, cross B: N = 87). Using the R (v4.1.2) package *dplyr* (v1.0.7), we then randomly selected 80 granulomas from each group for the analysis. To assess the quantity of fibrosis per sample, we trained HALO’s Area Quantification tool (v2.3.1) on the different trichrome dyes (blue, red, brown), while accounting for potential batch staining variability. This tool automatically detects pixels of an assigned color and calculates the total area stained with each dye.

### *Measurement of* S. mansoni *larval production and snail response to infection*

#### a. Number of viable LS and HS miracidia

To recover F3 schistosome eggs, we pooled livers of 10 mice infected with each parasite line (LS and HS) and crosses (A and B). We processed each of the four pool independently as described in (23). We then counted the number of viable miracidia hatched from each pool of eggs for each line. Briefly, we sampled four 20 μL aliquots of freshwater containing miracidia larvae, added 20 μL of 20X normal saline and counted the immobilized miracidia under a microscope. We estimated the number of miracidia/mL and the overall number of miracidia in the total volume of the miracidial solution. To obtain the number of miracidia/gram of liver, we divided the overall number of miracidia estimated by the exact weight of the livers processed for a given group of mice (either infected with LS or HS selected line).

We evaluated the reproductive fitness of each parasite line per cross by dividing the total number of miracidia by the total number of female worms recovered after mouse perfusion.

#### b. Cercarial production of LS and HS parasite lines in different snail populations

We measured cercarial production of LS and HS parasites. We infected two inbred snail populations to determine if the virulence of the selected lines varies with the snail host genotype. We used Bg26 snails (N= 240 exposed snails/selected lines/cross), and Bg121 snails (N= 144 exposed snails/selected lines/ cross). We exposed each snail to a single miracidium to allow examination of cercarial shedding from single parasite genotypes (16) (Figure 1).

Every week, from week 4 to 7 post-exposure, we assessed cercarial production of individual Bg26 and Bg121 snails infected with either F3 LS or HS selected parasites lines for each cross (16).

#### c. Snail physiological response to parasitic infection

We measured the impact of infection on the snail host by quantifying daily snail survival and physiological responses (laccase-like activity and hemoglobin rate in the hemolymph) at end point. During week 7, 3 days after the last cercarial shedding, we collected hemolymph (24) from all surviving snails infected with either LS or HS line for each cross. We measured both laccase-like activity (24) and hemoglobin rates in the hemolymph of each infected snail (4).

### Statistical analysis

All statistical analyzes and graphs were performed using R software (v4.1.2) (25). When data were not normally distributed (Shapiro test, p < 0.05), we compared results with non-parametric Kruskal-Wallis test followed by a pairwise Wilcoxon post-hoc test, or by pairwise comparison using Wilcoxon-Mann-Whitney test (two groups comparison). When data followed a normal distribution and had homogeneous variance (Bartlett test, *p* > 0.05), we used one–way ANOVA followed by Tukey HSD post-hoc test or a pairwise comparison Student *t*-test or Welsh *t*-test (two groups comparison). We used binomial tests to assess sex ratio differences. Variation in snail susceptibility to the different lines of *S. mansoni* parasites were analyzed using Chi-square tests. We performed snail survival analyses using log-rank tests (R survival package, v3.4-0) and correlation analysis with Pearson’s correlation test. *p*-values less than 0.05 were considered significant.

## RESULTS

### *Increased virulence of the high shedder* S. mansoni *to the rodent host*

Our HS and LS populations were founded from the two extremes of the F2 phenotypic distribution. These two lines have the same genetic background, but are expected to differ at QTL regions that underlie cercarial shedding (16). We can use these lines to examine how genes that determine cercarial shedding impact fitness characteristics and virulence of adult parasite in the vertebrate host.

We found no differences in growth rate of mice infected with the HS parasite line compared to the LS line or to the control group. All the mice gained weight over the time of the experiment (Figure 2A; Kruskal-Wallis test, KW = 119.81, df = 6, *p* < 2.2×10^−16^), independently of infection status.

**Figure 2:**
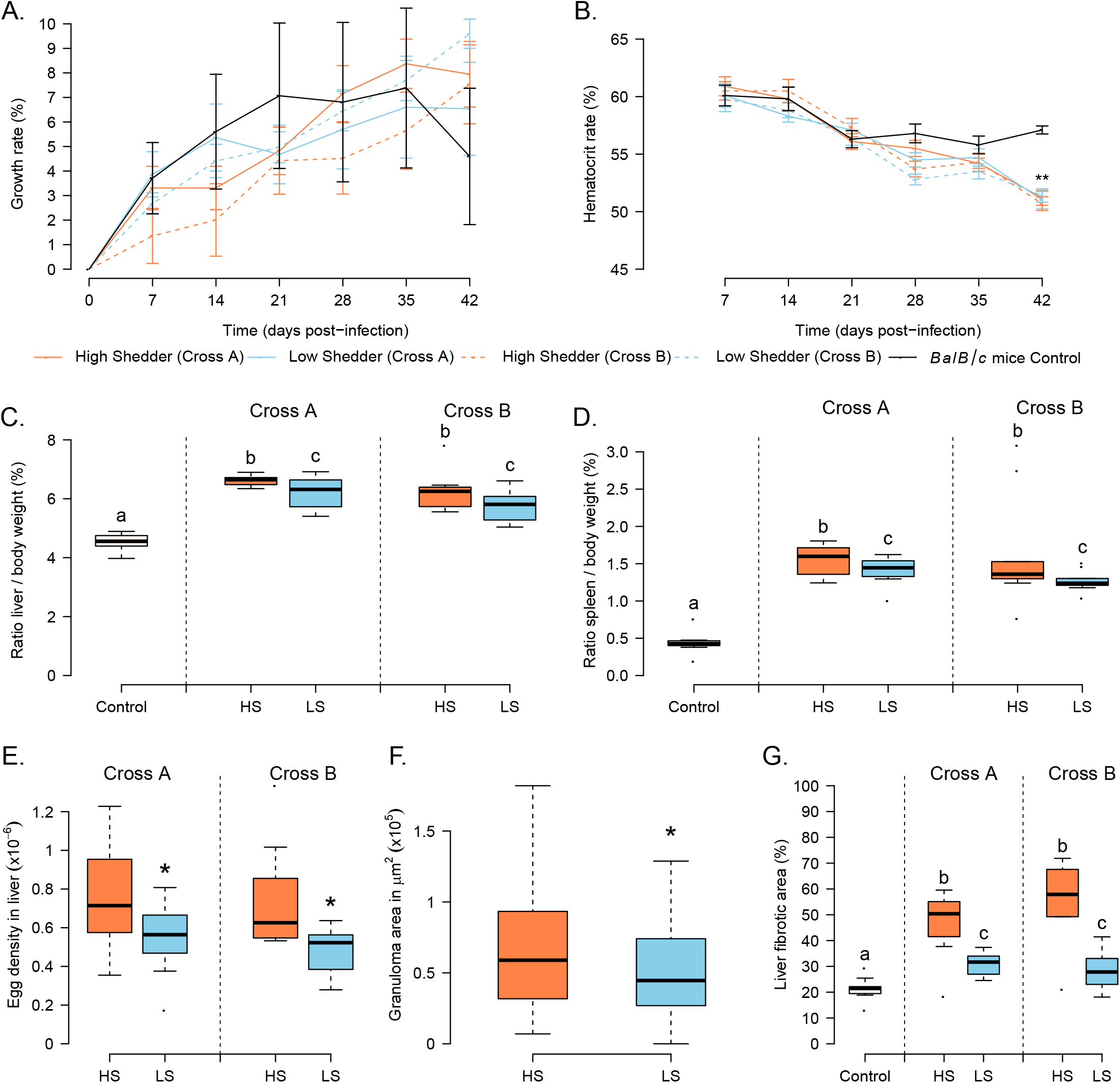
Differential impact of LS and HS parasites on rodent host life-history traits and pathology. LS and HS *S. mansoni* lines do not differentially affect the BALB/c growth rate compared to control group **(A)** or hematocrit rate **(B)** over the course of infection. However, mice infected with HS parasites exhibit greater hepatosplenomegaly compared to rodents infected with LS lines **(C)** Ratio liver/body weight (%) and **D**. Ratio spleen/body weight (%)]. Similarly, livers from HS infected mice exhibited higher egg density **(E)**, associated with the formation of bigger granulomas **(F)** and increased hepatic fibrosis **(G)** compared to livers from LS infected mice. Groups (parasite lines or control) not connected by the same letter are significantly different (post-hoc test). \**p* < 0.05; ** *p* ≤ 0.01; *** *p* ≤ 0.001.

As one symptom of schistosomiasis is anemia (26), we followed the hematocrit rate of each infected or control mouse over the course of the infection. The hematocrit rate of the infected mice is significantly lower compared to the control group (Figure 2B; Kruskal-Wallis test, KW = 11.66, df = 4, *p* = 0.020, followed by Pairwise Wilcoxon test). This is particularly striking at t+42 days post-infection (Pairwise Wilcoxon test, *p* = 0.0015), when all the worms are fully mature and are feeding on red blood cells (27, 28). However, there was no impact of HS or LS infections on hematocrit.

The ratio liver/body weight is higher in mice exposed with the selected HS parasite lines compared to the LS ones (Figure 2C; Kruskal-Wallis test, KW = 26.92, df = 2, *p* = 1.42×10^−6^, followed by Pairwise Wilcoxon test). This result was not associated with worm burden, as we found no differences between numbers of worms in HS and LS infected mice (Figure 3A; Cross A: Wilcoxon test, W = 55, *p* = 0.7331; Cross B: Welsh t-test, t= 1.5598, df=18, *p* = 0.1362). Similarly, we detected no differences in the sex ratio (number of males / number of females) of the recovered worms when comparing HS versus LS lines (Cross A: binomial test, HS line (sex ratio = 0.89): *p* = 0.3596, LS line (sex ratio = 0.82): *p* = 0.1221; Cross B: binomial test, HS line (sex ratio = 0.94): *p* = 0.6821, LS line (sex ratio = 1.1): *p* = 0.5107). This result suggests that HS lines produced more eggs, which become trapped in mouse liver causing increased hepatomegaly symptoms.

**Figure 3:**
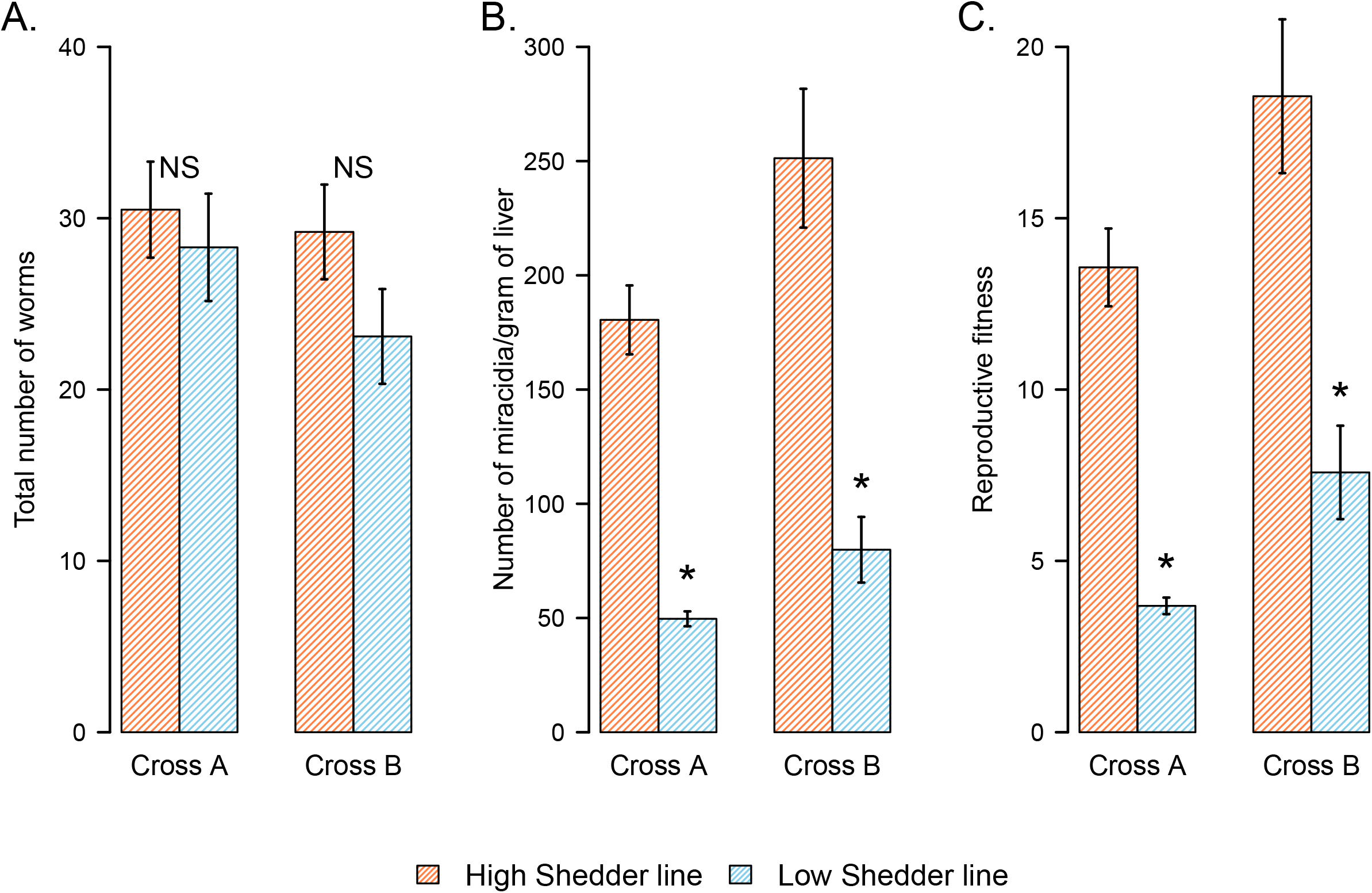
Increased reproductive fitness of HS parasite line in rodents. **(A)** The ability of cercarial larvae to infect its rodent definitive host is similar for LS and HS lines, in both genetic crosses, as a comparable number of worms were recovered from mice infected with HS and LS parasites after being exposed to the same number of cercariae. **(B)** In both crosses, the HS parasite line produced significantly more live miracidia (recovered from rodent infected livers) compared to the LS parasites, and therefore **(C)** HS reproductive fitness (total number of miracidia recovered/total number of female worms) is higher than in LS parasites. NS: No significant difference between the considered groups; \**p* < 0.05; ** *p* ≤ 0.01; *** *p* ≤ 0.001.

The histopathology analysis confirmed that HS infected mice had a higher egg density than LS infected mice (Figure 2E; Cross A: Welsh t-test, t = 2.2745, df = 15.705, *p* =0.0373; Cross B: Wilcoxon test, W = 83, *p* = 0.0115). Interestingly, HS lines also induced the formation of larger granulomas than LS lines (Figure 2F; Wilcoxon test, W = 15276, *p* = 0.0027). We found no correlation between the average granuloma size and the number of eggs in liver (Pearson’s correlation test, *p* = 0.0678, *r* = 0.29). As a result of egg entrapment and granulomatous response, livers from infected mice showed an increase in collagen deposition indicative of fibrosis compared to control mice (Figure 2G; Kruskal-Wallis test on the complete dataset, KW = 29.109, df = 4, *p* = 7.43×10^−6^, followed by Pairwise Wilcoxon test; Cross A: KW = 17.809, df = 2, *p* = 0.00013; Cross B: KW = 17.05, df = 2, *p* = 0.00019). Notably, we found that fibrotic area was significantly enlarged in livers infected with HS selected lines compared to livers infected with LS lines.

The ratio spleen/body weight (expressed in %) is also higher in mice infected with HS parasites (Figure 2D; Kruskal-wallis test, KW = 26.037, df = 2, *p* = 2.21×10^−6^, followed by Pairwise Wilcoxon test) suggesting a stronger immune response.

### Increased reproductive fitness of HS parasite line in rodents

We observed significantly more viable miracidia isolated from the liver of mice infected with the HS parasite compared to the LS parasite for both crosses (Figure 3B; Cross A: Wilcoxon test, *p* = 0.0265; Cross B: Wilcoxon test, *p* = 0.0284). The higher number of viable miracidia produced by HS parasite lines is not the consequence of a higher worm burden (Figure 3A), but is explained by a significantly higher reproductive fitness of the HS compared to LS female worms (Figure 3C; Cross A: Wilcoxon test, *p* = 0.02652; Cross B: Wilcoxon test, *p* = 0.02843).

### Fitness and transmission parameters of LS and HS parasites in different snail lines

When we examined the transmission phenotype of the F3 progeny from these selected lines, we found that the HS parasites produced significantly more cercariae than the LS parasites regardless of the genetic background of the snail (Bg26 or Bg121) (Figure 4A-B).

**Figure 4:**
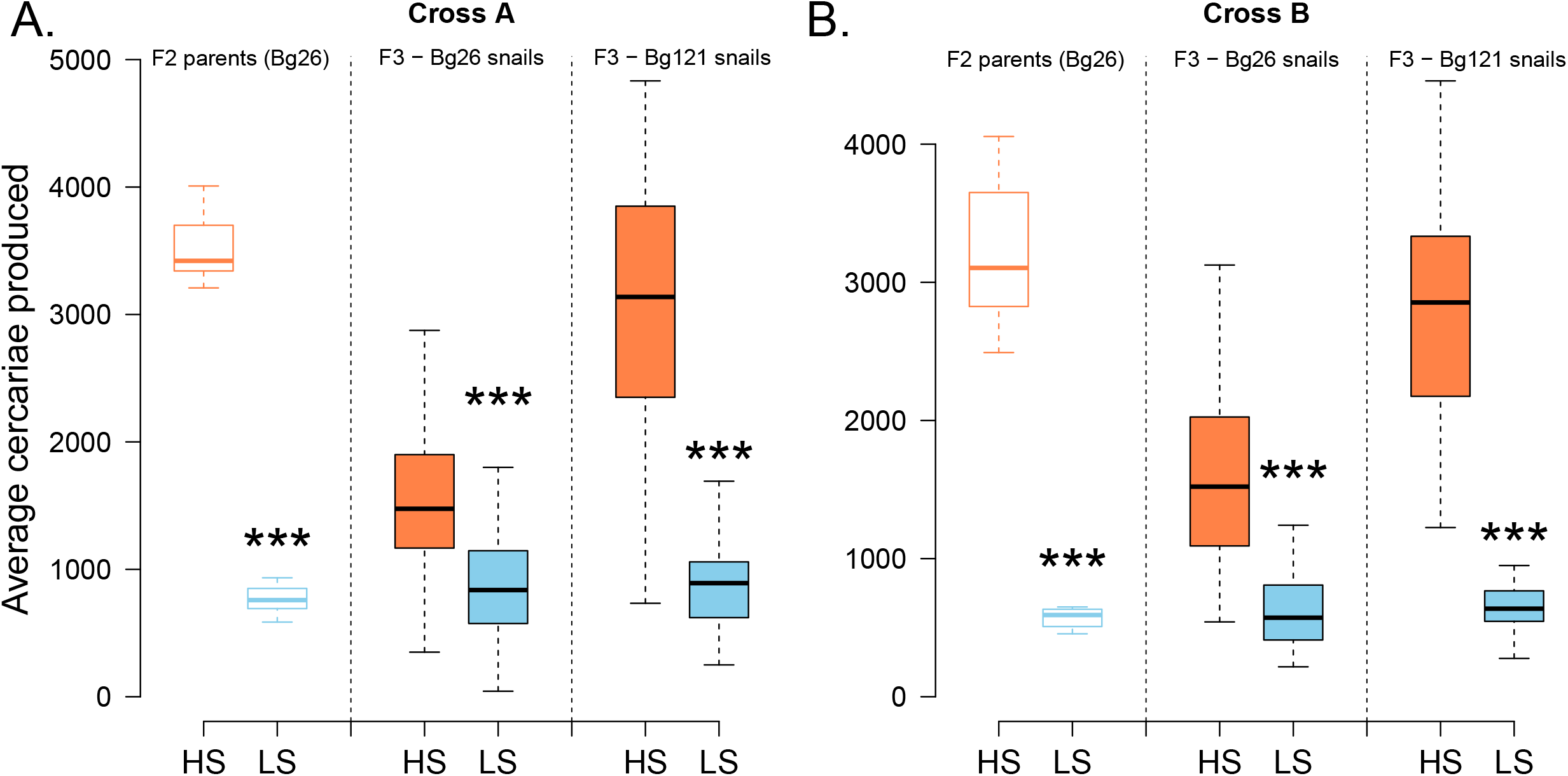
Differences between LS and HS parasite life-history and transmission traits in different snail lines. HS and LS lines were founded by selecting F2 parasites exhibiting extremely low and extremely high shedding phenotypes, for both genetic crosses (A and B) between SmBRE-LS and SmLE-HS. **(A-B: F2 parents)** For both crosses, F2 LS parents (in Bg26 snails) produced fewer cercariae (average over 4 weeks, one shedding/week) compared to F2 HS parents (in Bg26 snails) (Cross A: Wilcoxon test, W=100, *p* = 1.083×10^−5^; Cross B: Welsh t-test, t= 10.933, df = 35.838, *p* = 1.334×10^−8^). Similarly, for both crosses **(A-B: F3)** and snail genetic background (Bg26 or Bg121), F3 progeny HS parasites produced significantly more cercariae (average over 4 weeks, one shedding/week) than F3 LS parasites (Cross A, F3-Bg26 snails: Welsh t-test: t = 5.8502, df = 80.313, *p* = 1.019×10^−7^, F3-Bg121 snails: Welsh t-test: t = 10.933, df = 35.838, *p* = 5.76×10^−13^; Cross B, F3-Bg26 snails: Wilcoxon test: W = 3084, *p* = 8.88×10^−15^, F3-Bg121 snails: Welsh t-test: t = 15.557, df = 39.28, *p* < 2.2×10^−16^). \**p* < 0.05; ** *p* ≤ 0.01; *** p≤ 0.001.

Bg121 snails shed significantly more cercariae than Bg26 when infected with HS parasites (Figure 4A-B; Cross A, Kruskal-Wallis test: KW = 32.769, df = 1, *p* = 1.038×10^−8^; Cross B, Kruskal-Wallis test: KW = 34.802, df = 1, *p* = 3.65×10^−9^). However, Bg26 and Bg121 snails infected with LS lines showed no difference in cercarial production in either cross (Figure 4A-B; Cross A, Kruskal-Wallis test: KW = 0.0510, df = 1, *p* = 0.821; Cross B, Kruskal-Wallis test: KW = 0.4625, df = 1, *p* = 0.4964).

Interestingly, we found no impact of snail population (Bg26 vs Bg121) or parasite line (LS vs HS) on susceptibility to infection (Bg26 vs. Bg121 infected with HS - Cross A - χ2 test : χ = 0.003, df = 1, *p* = 0.9535; Cross B: χ = 2.9175, df = 1, *p* = 0.087; Bg26 vs. Bg121 infected with LS - Cross A:, χ = 0.017, df = 1, *p* = 0.8951; Cross B: χ = 2.0707, df = 1, *p* = 0.1502; Cross A HS vs. LS in Bg26 - χ2 test : χ = 0.0634, df = 1, *p* = 0.801; HS vs. LS in Bg121, χ = 0.1059, df = 1, *p* = 0.7448; Cross B HS vs. LS in Bg26 - χ2 test, χ = 2.5255, df = 1, *p* = 0.112; HS vs. LS in Bg121, χ = 3.4526, df = 1, *p* = 0.063).

The striking differences in miracidia and cercariae production between HS and LS selected lines of *S. mansoni* parasites showed limited dependence on host type (rodent or snail) or mollusk genetic background (Bg26 or Bg121) but were strongly linked to parasite genetics (16).

### Major impact of HS parasite on the snail host physiology

We found no significant difference in survival of snails (Bg26 and Bg121) infected with HS and LS parasites (Figure 5A, Cross A: LogRank test, χ = 5.2, df = 3, *p* = 0.2; Cross B: LogRank test, χ = 4.5, df = 3, *p* = 0.2). However, we found a significant differential impact of these two selected lines on snail host physiology. Laccase-like activity (Figure 5B) and hemoglobin rate (Figure 5C) measured in the snail hemolymph, 7.5 weeks post-infection, provide good proxies to evaluate snail health and the impact of schistosome infection (16,23,29). Both parameters show strong reduction in snails infected with HS selected lines, independently of the cross or the snail population (Figure 5B and Figure 5C).

**Figure 5:**
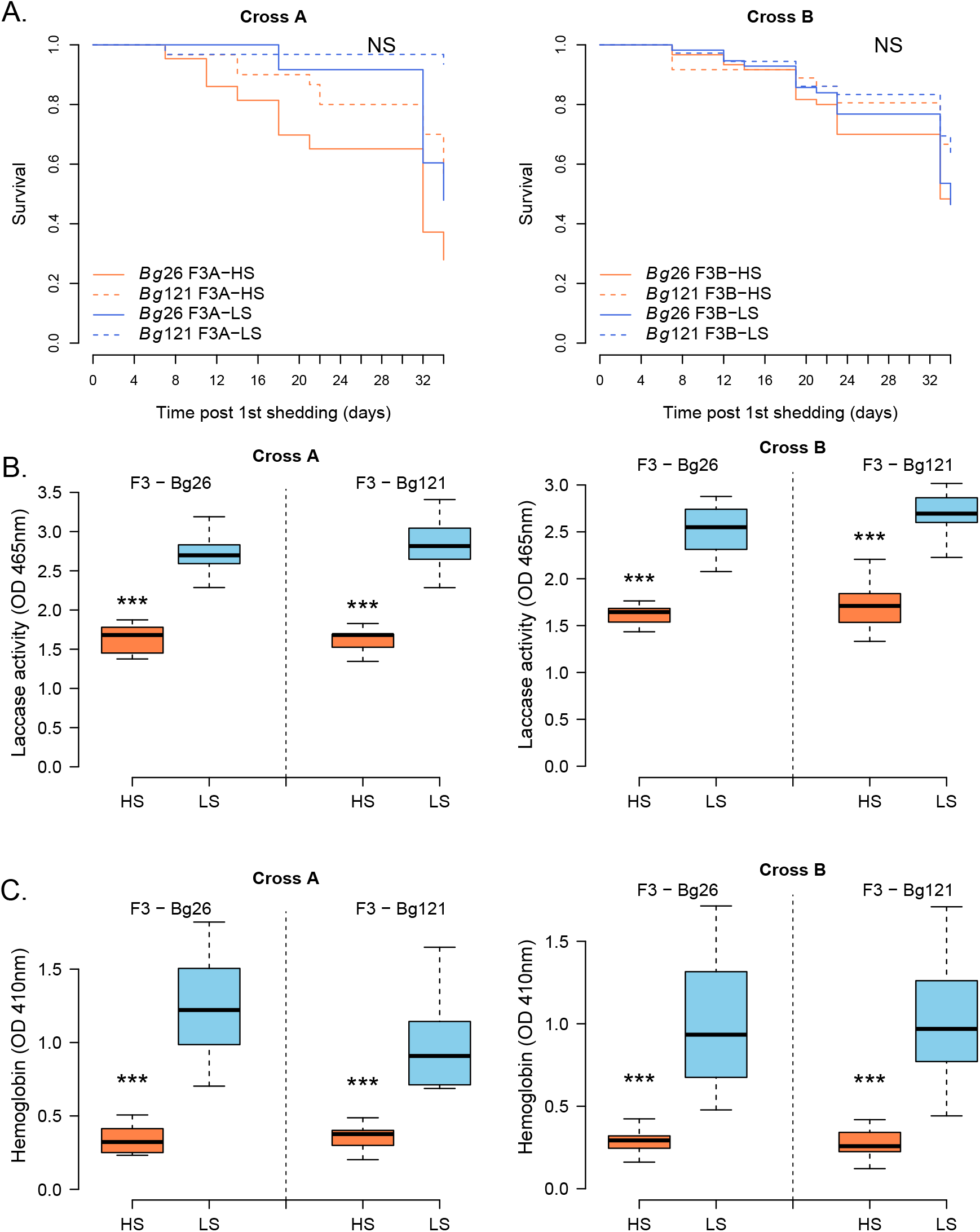
Differential impact of HS and LS parasites on snail host life-history traits and physiology. **(A)** We did not detect significant differences in the survival of the infected snails between populations (Bg26 or Bg121) or between infection groups (HS or LS schistosome parasites selected lines). **(B-C)** There was a strong impact of the infection group on the snail physiological parameters. **(B)** laccase-like activity (Welsh t-test: Cross A: F3-Bg26 infected with HS vs. LS, t = −11.126, df = 16.846, *p* = 3.50×10^−9^; F3-Bg121 infected with HS vs. LS, t = −9.3019, df = 12.838, *p* = 4.57×10^−7^; Cross B: F3-Bg26 infected with HS vs. LS, t = −11.329, df = 13.79, *p* = 2.28×10^−8^; F3-Bg121 infected with HS vs. LS, t = −9.6534, df = 25.15, *p* = 6.11×10^−11^) and **(C)** hemoglobin rate (Welsh t-test: Cross A: F3-Bg26 infected with HS vs. LS, t = - 7.8325, df = 12.101, *p* = 4.429×10^−6^; F3-Bg121 infected with HS vs. LS, t = −6.4313, df = 12.051, *p* = 3.185×10^−5^; Cross B: F3-Bg26 infected with HS vs. LS, t = −6.0988, df = 11.636, *p* = 6.099×10^−5^; F3-Bg121 infected with HS vs. LS, t = −7.3274, df = 14.353, *p* = 3.238×10^−6^). Snails infected with HS schistosome parasites lines consistently exhibit lower laccase activity and hemoglobin rates compared to those infected with LS parasite lines. NS: No significant difference between the considered groups; \**p* < 0.05; \*\**p*≤0.01; \*\*\**p*≤ 0.001.

Both laccase-like activity and hemoglobin rate were negatively correlated with F3 cercarial production in both snail populations (Pearson’s correlation tests - Cross A - Laccase-like activity: *p* = 8.052×10^−6^, r = −0.64; Hemoglobin rate: *p* = 5.119×10^−5^, r = −0.60; Cross B: Laccase-like activity: *p* = 1.91×10^−8^, r = −0.68; Hemoglobin rate: *p* = 9.783×10^−7^, r = −0.61). We observed a strong positive correlation between these two physiological parameters (Pearson’s correlation tests - Cross A: *p* = 3.987×10^−13^, r = 0.87; Cross B: *p* < 2.2×10^−16^, r = 0.86) as reported previously (16, 23).

## DISCUSSION

### Differences between LS and HS parasite virulence and fitness in the rodent host

The phenotypic characterization of LS and HS parasite lines in the rodent and snail host clearly showed no evidence for trade-offs in parasite fitness between intermediate and definitive hosts Figure 6. When comparing life history traits and virulence of LS and HS parasites in the rodent host, we observed that LS parasites show low virulence in mice with fewer eggs in the liver, smaller granulomas with less fibrosis of the hepatic tissue. These parasites produced less viable miracidia larvae (Figure 6). On the other hand, HS *S. mansoni* exhibits high virulence in mice with more eggs in the liver, bigger granulomas associated with significantly more fibrosis of the hepatic tissue. These parasites produced more viable miracidia larvae (Figure 6).

**Figure 6:**
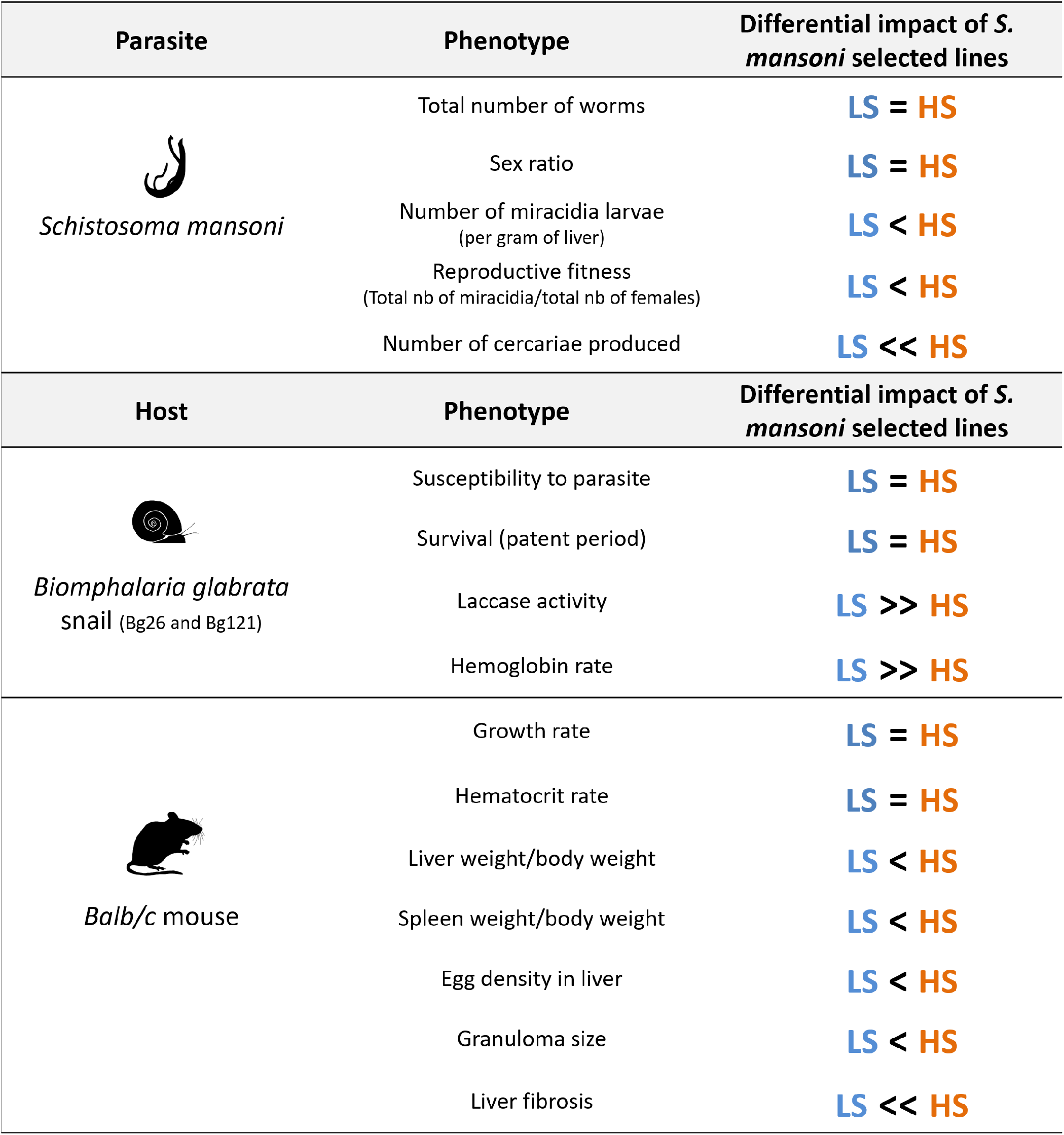
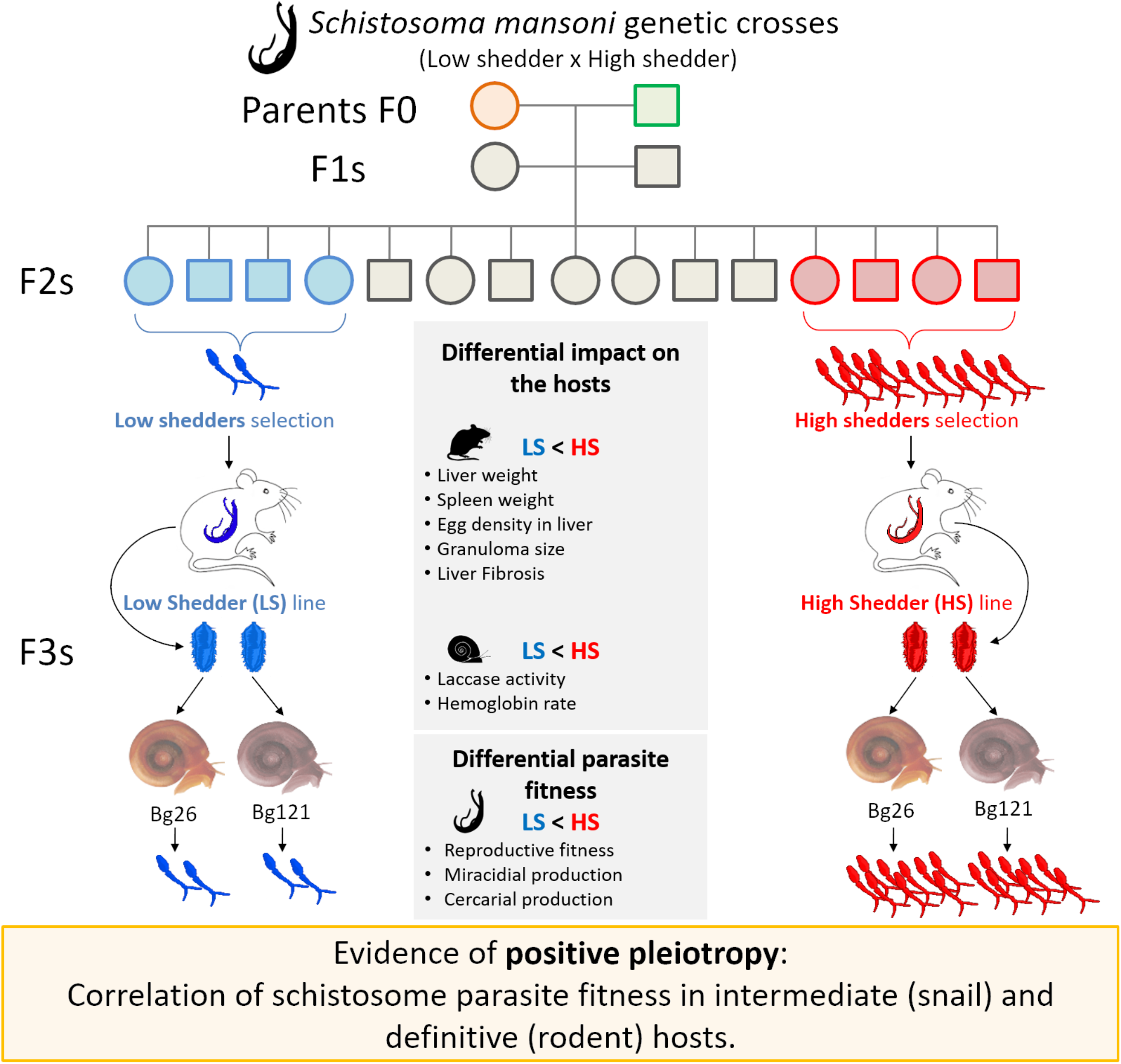
Differential impact of schistosome selected lines on the phenotype of their intermediate and definitive hosts. Summary table of all the phenotypes measured for the selected HS and LS *S. mansoni* lines and the differential impact of these selected lines on the phenotype of their snail intermediate hosts and rodent definitive host.

We also demonstrated that the greater virulence of HS parasites to rodent host does not result in greater infectivity to mice; we showed no difference in the number of worms recovered between the two selected lines (HS and LS) for each cross (Figure 6). However, we noticed a significantly higher reproductive fitness of HS compared to LS parasites, with more viable miracidia/worm (Figure 6).

### A signature of positive pleiotropy for fitness in our schistosome model

Using experimental selection for parasite infection intensity in one African and one Puerto-Rican (SmPR) population of *S. mansoni*, Gower and Webster (5) showed opposite patterns of fitness in snail intermediate and vertebrate definitive host (antagonistic pleiotropy). Similar conclusions had been drawn by Davies et al. (30), who demonstrated a trade-off in parasite (SmPR) reproductive success in the intermediate and definitive hosts. However, in the present study, we found that schistosome fitness was correlated in intermediate and definitive host. These results reveal no fitness trade-off and the presence of positive pleiotropy in our schistosome-snail-rodent model.

Differences between our conclusions and those presented by Davies *et al*. (30) and Gower and Webster (5) could be explained by multiple factors, including i) the different populations of schistosome parasites investigated (SmPR in Davies *et al*. (30) and Gower and Webster (5) versus the progeny of crosses between SmLE and SmBRE *S. mansoni* populations in the present study); ii) differing dose of miracidia larvae used to infect snails (six miracidia/snail in Davies *et al*. (30) and seven miracidia/snail in Gower and Webster (5) versus one miracidium/snail in our study); iii) differences in the selection process of the low and high shedders (based on a single shedding in Gower and Webster (5) versus on the average of four weekly cercarial shedding in the our study); iv) the genetic background of the intermediate snails and definitive mice hosts (CBA/CA mice in Davies *et al*. (30) and Gower and Webster (5) versus BALB/c mice in this study). Murine genetic background is known to affect the fecundity of *S. mansoni* worms (31).

Interestingly, we also demonstrated that HS parasites are not only characterized by higher reproductive fitness in their rodent hosts and increased splenomegaly linked to a severe immune response, but their eggs trigger a stronger immune reaction in the mouse liver, leading to bigger granulomas. Granulomas are a result of inflammation around the eggs, due to the strong T-helper 2 immune response that they induce (32). However, the size of the granulomas depends on the *S. mansoni* parasite population (33). As most schistosomiasis disease symptoms are caused by granuloma formation, it would be interesting to investigate the composition of eggshell proteins (32) produced by LS and HS *S. mansoni* parasites.

### Why do we see positive pleiotropy in HS and LS schistosome parasites?

Genetic variants that result in low fitness in snails and the adult stages of the parasites seem to be disadvantageous and should be selected against. We know the QTLs that underly cercarial shedding number (16). The alleles determining low shedding number are all inherited from the SmBRE parent. This paper demonstrates that these same genome regions also negatively impact fitness and virulence of adult parasites in the vertebrate host. What might explain the existence of variants that have negative consequences in both invertebrate and vertebrate host? Inbreeding depression and genetic drift could be the causes of low transmission stage production in LS parasites (and SmBRE) if several detrimental loci affecting both cercariae and miracidia production are fixed. These could explain the relatively poor fitness of LS compared to HS parasites. The QTLs underlying low cercarial shedding (16) contain deleterious alleles at loci impacting transmission. However, these loci do not affect at all the compatibility between LS parasites and their hosts, as LS is perfectly able to infect both snails and rodents at the same rate as HS parasites. To investigate the “inbreeding hypothesis”, we predict that the SmLE population shows diminished variation relative to other laboratory populations of schistosome parasites.

There are several caveats to our interpretation of these results. First, the measures of fitness that we have used are incomplete. Our present study examined hepatic pathology caused by *S. mansoni* infection and demonstrated that LS selected parasites induced less hepatic fibrosis, associated with less and smaller granulomas present in the liver compared to HS parasites. However, we did not investigate the egg burden and viable miracidia present in the intestinal epithelia and in mouse feces, the natural transmission route for *S. mansoni*. Therefore, the apparent “low reproductive fitness” of LS parasites could be biased by the egg tissue tropism (liver versus intestine). LS parasites and their parent SmBRE may be more fit than initially thought: while HS parasites (and SmLE) are fully adapted to a laboratory life, where schistosome parasites are transmitted through eggs collected from mammal liver (a natural dead-end for *S. mansoni*), LS parasites (and SmBRE) may have retained a more natural egg tropism, where most eggs pass through the intestinal wall and are excreted in mammal feces.

Second, mice are laboratory hosts for schistosomes. While we have observed positive pleiotropy using the mouse host, it is conceivable that LS parasites might show high fitness in the natural human host. However, we think this is unlikely as we observed low fitness and pathology of SmBRE in hamster hosts and in different mouse lines (Jutzeler *et al*. unpublished).

Third, further work may reveal that parasite alleles associated with low shedding of cercariae from Bg26 and Bg121, result in high shedding in additional snail populations. Our demonstration that shedding phenotype is comparable in two different snail populations argues against this.

## CONCLUSIONS

In this study, we showed positive pleiotropy and absence of trade-off in transmission strategy and virulence from *S. mansoni* LS and HS, for both snail intermediate and rodent definitive hosts. We demonstrated that genetic loci (or co-segregating genes) involved in cercarial production influence adult worms’ reproductive fitness and egg production. However, while it is clear that high cercarial production in snail hosts is associated with an increase in pathogenicity in rodent hosts (measured as hepatosplenomegaly and liver fibrosis), it is still unclear whether the low pathogenicity exhibited by the LS is due to a true low reproductive fitness of the worms or to differing tissue tropism of the eggs. More generally, our study demonstrates how genetic variants may determine parasites phenotypes at different stages of the lifecycle.

## DECLARATIONS

### AVAILABILITY OF DATA AND MATERIALS

All the phenotype datasets and the R scripts used for data analysis, to generate the Figures (Figures 2 to 5), and performe statistical tests are available in the Zenodo repository at https://doi.org/10.5281/zenodo.7311858. All the histopathology datasets analyzed in this manuscript are available in the Zenodo repository at [in process].

### COMPETING INTERESTS

The authors declare that they have no competing interests.

### FUNDINGS

This research was supported by a Cowles fellowship (WL) from Texas Biomedical Research Institute (13-1328.021), by a Graduate Research in Immunology Program training grant NIH T32 AI138944 (KJ), and NIH R01 AI133749 (TJCA), and was conducted in facilities constructed with support from Research Facilities Improvement Program grant C06 RR013556 from the National Center for Research Resources. SNPRC research at Texas Biomedical Research Institute is supported by grant P51 OD011133 from the Office of Research Infrastructure Programs, NIH.

### AUTHORS’ CONTRIBUTIONS

WL, FDC and TJCA designed the genetic crosses of *Schistosoma mansoni* parasites and the selection experiments. WL and FDC performed all the experiments (schistosome genetic crosses and selections, measurement of all the phenotypes in parasites and hosts – snails and rodents). WL performed all the data analyses, KJ performed the analysis of rodent liver histopathology. WL and TJCA drafted the manuscript. All authors read and approved the final manuscript.

## ACKNOWLEDGEMENTS

We thank Michael S. Blouin (Oregon State University) for providing the Bg26 and Bg121 snail lines. We thank the Vivarium of the Southwest National Primate Research Center (SNPRC) for providing rodent care. We also thank Renee Escalona, Colin Chuba and Jesse Martinez from the Hixon Pathology core lab (Texas Biomedical Research Institute) for performing all the histopathology preparations and staining of mouse livers and Dr. Vinay Shivanna for providing access to the HALO software for analyzing the histopathology slides.

## Notes

https://doi.org/10.5281/zenodo.7311858

### Competing Interest Statement

The authors have declared no competing interest.

